# Taking Fright: The Decline of Australian Pied Oystercatchers *Haematopus longirostris* at South Ballina Beach, New South Wales

**DOI:** 10.1101/2020.06.09.141796

**Authors:** Stephen Totterman

## Abstract

This study reviewed data from the Richmond River Area Pied Oystercatcher Protection Program 1997–2013 and Richmond River Area Shorebird Protection Program 2014–2018, on the far north coast of the State of New South Wales (NSW), Australia. The Australian Pied Oystercatcher *Haematopus longirostris* breeding population size for South Ballina Beach has decreased from 15–16 pairs in 1994–1996 to 7–9 pairs in 2016–2018. This is despite control of the European Red Fox *Vulpes vulpes* successfully reducing depredation of eggs and chicks and > 207 oystercatchers fledging from beaches in the Richmond River area in 1997–2018. The negative trend for South Ballina is contrasted with the positive trend for Bombing Range Beach, where the population size has increased from 4–5 pairs in 2002–2004 to 8–9 pairs in 2016–2018. Vehicle-based recreation has increased at South Ballina in the past two decades versus Bombing Range has remained closed to the public. It is proposed that human recreation disturbance is preventing recruitment into the South Ballina oystercatcher breeding population. Without strong protection of habitat from such disturbance, the NSW oystercatcher breeding population size is predicted to continue to decrease.

## INTRODUCTION

The Australian Pied Oystercatcher *Haematopus longirostris* inhabits estuaries and ocean beaches along the Australian coast, with major populations in the southern states of Tasmania, South Australia and Victoria. The species is non-migratory and many breeding pairs remain on their territories throughout the year. It is also long-lived and breeding population sizes are usually limited by territorial behaviour and not by breeding success. Delayed age of first breeding is normal, resulting in a surplus of non-territorial adults in populations (see the conservation assessment by Taylor *et al.* 2014).

The Australian Pied Oystercatcher has been assessed as Endangered in New South Wales (NSW) because the estimated population size is low and projected or continuing to decrease. Proposed conservation threats for the species include habitat loss, human recreation disturbance, a reduction in food resources and low breeding success, often because of depredation of eggs and chicks by the European Red Fox *Vulpes vulpes* (NSW Scientific Committee 2010). The NSW Department of Planning, Industry and Environment (DPIE) (2019a, 2019b) has estimated that there are 200 oystercatcher breeding pairs in NSW and Totterman (2020) estimated that < 54 of those are ‘beach-residents’, *i.e.* birds that forage, roost and spend most of the time on the beach. The Richmond River Area, between the Richmond and Clarence Rivers, supports the largest numbers of beach-resident oystercatchers in NSW, with 22–23 breeding pairs in 2016–2018 (this study).

Since 1997, the primary conservation action for oystercatchers in the Richmond River Area has been fox control to reduce predation of eggs and chicks (Wellman *et al.* 2000). This action has received legislative support from the NSW Threatened Species Conservation Act 1995 and its successor, the NSW Biodiversity Conservation Act 2016 (NSW Government 2016), Sch. 4 of which lists Predation by the Red Fox as a Key Threatening Process (in general and not only for oystercatchers). This action has received financial and management support from the NSW Government in successive Threat Abatement Plans for predation by the Red Fox (NSW National Parks & Wildlife Service 2001, NSW Office of Environment & Heritage 2010). The number of priority sites for the Australian Pied Oystercatcher has increased from 13 in the first plan to 19 in the second. This breeding success conservation strategy assumes that sufficient numbers of oystercatcher fledglings will: 1) survive to become adults; 2) stay in or return to NSW, and; 3) be successful in acquiring a breeding territory.

Oystercatchers are specialist predators of bivalves and Owner and Rohweder (2003) reported that oystercatchers on far north coast beaches of NSW are associated with high abundance of the surf clam *Donax deltoides*, commonly known as the ‘pipi’. Pipi stocks ‘crashed’ between 2003–2009 and the associated negative fluctuation in oystercatcher counts was interpreted by Harrison (2009) and then the NSW Scientific Committee (2010) as a permanent reduction in the population size. Harrison (2009) also proposed that low pipi abundance can result in low oystercatcher breeding success. Totterman (2018) reported that oystercatcher counts for South Ballina Beach did increase when pipi stocks recovered *(i.e.* a numerical predator-prey response). However, Totterman (2020) reported that pipi abundance was a poor predictor for oystercatcher abundance on other beaches in NSW.

Birdlife Australia have stated that the greatest conservation threat to beach nesting birds is disturbance from people visiting the beach (Birdlife Australia 2018). Bird-habitat models for beach-resident Australian Pied Oystercatchers in Totterman (2020) included negative responses to urbanisation and pedestrian access density. Fisher *et al.* (1998) proposed that vehicle-based tourism on Fraser Island, Queensland, could negatively impact beach nesting bird populations via disturbance to their habitat. Harrison (2009) added that vehicle strikes are an additional source of mortality for shorebirds. Totterman (2020) noted that oystercatcher counts on 75 Mile Beach, Fraser Island, have decreased since Fisher *et al.* (1998) counted an average of 15 birds between Hook Point and Indian Head to a mean count of three in 2016.

Ferraro and Pattanayak (2006) noted that biodiversity conservation has too frequently been guided by intuition and anecdote and they called for more rigorous evaluation of conservation investments. This study applied impact evaluation methods to 25 years of breeding results for beach-resident oystercatchers in the Richmond River Area, looking at changes in breeding population sizes and variation in breeding success among sites in relation to fox control, pipi abundance and human recreation disturbance. Several questions were asked of the data: has the oystercatcher breeding population size increased following higher breeding success; is the breeding population size correlated with pipi abundance; is breeding success correlated with pipi abundance; has the breeding population size decreased at sites with more intense human recreation, and; is breeding success lower at sites with more intense human recreation?

## METHODS

### Study sites

The five sites in this study were defined by natural breaks in the beach habitat (*e.g.* headlands and rivers) and different management histories (Table 1, Figure 1).

**Figure 1.**
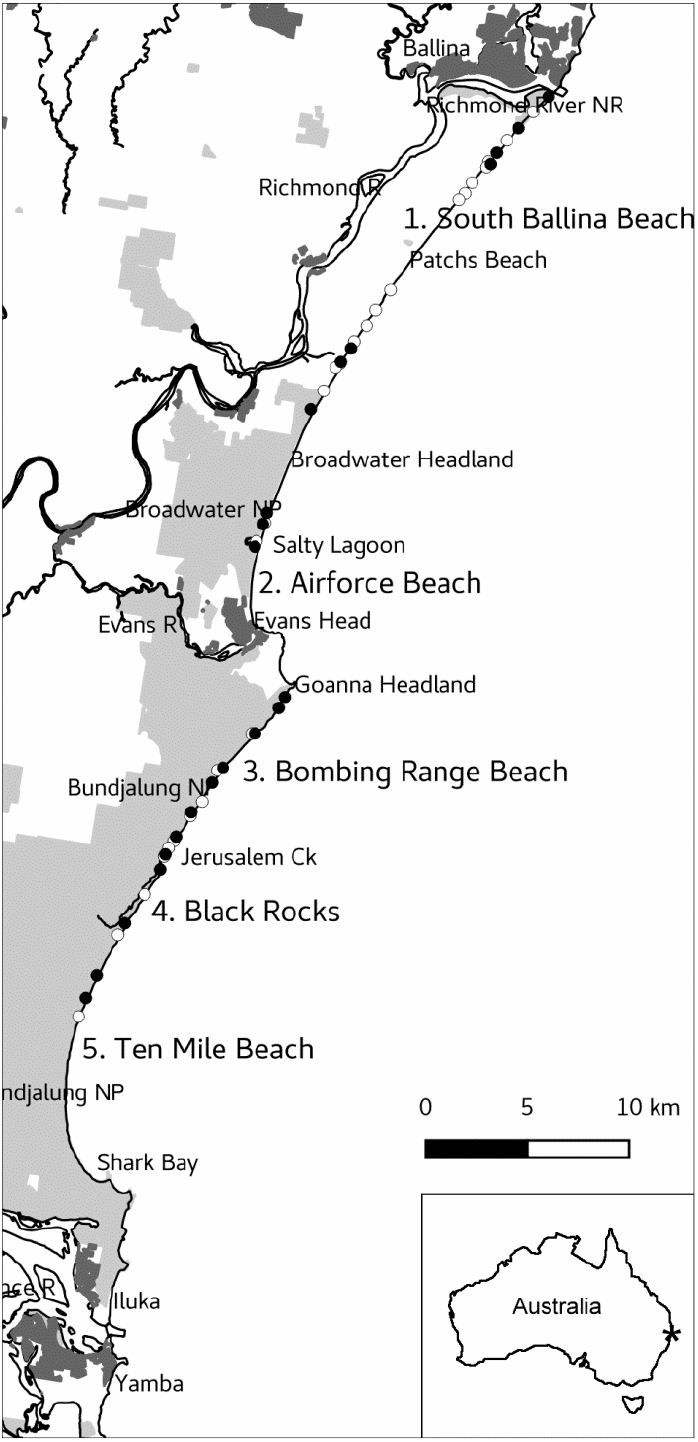
Map of the Richmond River Area with beach-nesting oystercatcher sites numbered 1–5 (Table 1). Light grey areas are National Parks and Nature Reserves. Dark-grey areas are urban. White-filled circles are 2005 nest locations (which was the first year where all sites were counted; *n* = 26). Black-filled circles are 2018 nest locations (*n* = 22). The first nest is plotted for pairs that made repeated attempts within a breeding season. The inset map shows the location of the Richmond River Area in Australia.

**Table 1.** Beach-nesting oystercatcher sites in the Richmond River area of NSW (Figure 1). Sites are defined by breaks in the beach habitat and different management histories. Data are from Totterman (2020) except that Black Rocks was not sampled in that study. Protected length includes adjacent Nature Reserves and National Parks (NP). Four wheel drive (4WD) length includes zones where beach driving was allowed. Dogs length includes zones where domestic dogs were allowed. Only public pedestrian access tracks were counted. The Black Rocks site adjoins a campground and walking track along Jerusalem Creek with many informal tracks to the beach. There is one track across the Goanna Headland isthmus to the northern end of Bombing Range Beach even though signs advise that public access is prohibited. Beach front human population is the sum of Australian Population Grid 2011 (Australian Bureau of Statistics 2011) 1 x 1 km cells intersected by the coastline.

| Site | N–S end | Total<br>(km) | Protected<br>(km) | Urban<br>(km) | 4WD<br>(km) | Dogs<br>(km) | Pedestrian<br>access | Camping<br>& c’van<br>parks | Human<br>pop. |
| --- | --- | --- | --- | --- | --- | --- | --- | --- | --- |
| 1. South Ballina | Richmond R. to Broadwater Hd.<br>(28.876–29.040°S) | 23.0 | 6.0 | 0.2 | 23.0 | 1.0 | 17 | 1 | 63 |
| 2. Airforce Beach | ‘Coffee rocks’ to Evans River<br>(29.060–29.113°S) | 5.8 | 4.3 | 1.3 | 4.6 | 1.3 | 4 | 1 | 173 |
| 3. Bombing Range | Goanna Hd. to Jerusalem Creek<br>(29.140–29.216°S) | 10.5 | 10.5 | 0.0 | 0.0 | 0.0 | 1 | 0 | 0 |
| 4. Black Rocks | Jerusalem Creek to Black Rocks<br>(29.216–29.254°S) | 4.8 | 4.8 | 0.0 | 0.0 | 0.0 |  | 1 | 0 |
| 5. Ten Mile | Black Rocks to Shark Bay<br>(29.254–29.358°S) | 12.5 | 12.5 | 0.0 | 12.5 | 0.0 | 2 | 0 | 0 |

South Ballina was defined as the 23 km long rural beach between the Richmond River and Broadwater Headland. A 2.3 km long ‘coffee rock’ (indurated sand) interval separates South Ballina from Airforce Beach. The N 2.5 km of South Ballina is part of Richmond River Nature Reserve, the S 3.5 km adjoins Broadwater National Park and the remaining 17 km adjoins a narrow Crown Reserve, backed by private properties. Beach driving was freely allowed S of the Richmond River Nature Reserve until April 2021.

Airforce was defined as the 5.8 km long beach N of the Evans River. This site includes Salty Lagoon (Salty Lakes), an intermittently closed and open lake. The N 4.3 km of Airforce adjoins Broadwater National Park. Beach driving was freely allowed N of the Evans Head township.

Bombing Range, Black Rocks and Ten Mile are contiguous sites adjoining Bundjalung National Park but have been managed differently (Table 1). Bombing Range was defined as the 10.5 km long beach between Goanna Headland (N) and Jerusalem Creek (S). Public access to Bombing Range was prohibited because it adjoins the Royal Australian Air Force Evans Head Air Weapons Range. The 4.8 km long Black Rocks (indurated sand) interval, where there was a campground, separates Bombing Range from the 12.5 km long S end of Ten Mile Beach. Beach driving was freely allowed N of Shark Bay to Black Rocks.

South Ballina, ‘Broadwater Beach’ (which was defined for management purposes as Airforce Beach N to Boundary Creek, at the S end of South Ballina) and Bombing Range are three Key Management Sites for the Australian Pied Oystercatcher in NSW (NSW DPIE 2019b).

To check if site definition bias affected the results, analyses were repeated for the aggregate sites ‘greater South Ballina Beach’ (South Ballina and Airforce) and ‘greater Ten Mile Beach’ (Bombing Range, Black Rocks and Ten Mile).

### Data sources

This study reviewed data from the Richmond River Area Pied Oystercatcher Protection Program 1997–2013 and Richmond River Area Shorebird Protection Program 2014–2018 (Appendix). Included are results from 1994–1996, preceding the start of fox control, and from 1997–2000, when there were no formal reports (Wellman *et al.* 2000). The conservation program was extended to include Airforce in 2001 and Bombing Range in 2002. Following the 2003–2009 pipi crash, the program was cancelled in 2011 and did not completely resume until 2016 (Table 2). The program was again cancelled in 2019, ostensibly because of the 2019 ‘Black Summer’bushfire crisis (Anon., pers. comm.).

**Table 2.**
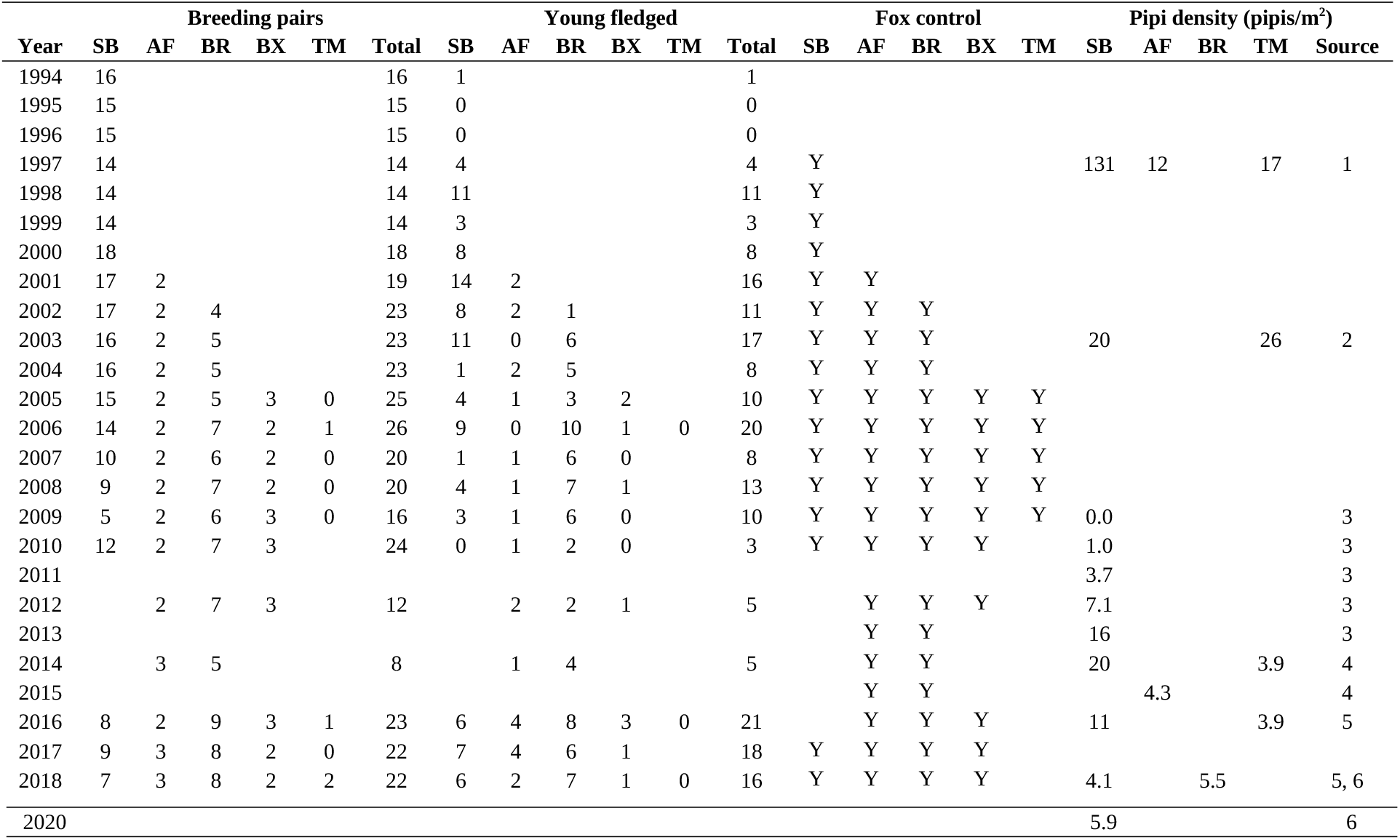
Oystercatcher breeding and pipi abundance results for South Ballina (SB), Airforce (AF), Bombing Range (BR), Black Rocks (BX) and Ten Mile (TM). Breeding data sources were: 1) Wellman *et al.* (2000) to 1999, and; 2) unpub. reports to the NSW DPIE thereafter (Appendix). Pipi data sources were: 1) Owner and Rohweder (2003); 2) Harrison (2009); 3) Totterman (2018); 4) Totterman (2019a); 5) Totterman (2020), and; 6) this study. Harrison (2009) did measure pipi abundance in 2005 but did not report those results for individual beaches. Blank results indicate no data including unsatisfactory oystercatcher monitoring in 2013 and 2015. Ten Mile was not surveyed in 2010 and 2011, however zero breeding pairs was assumed for those years to increase the Greater Ten Mile sample size. There was some fox control for South Ballina in 2011–2016 but it did not include the long Crown Reserve section. There was some fox control inland from Ten Mile Beach in 2010–2018 but not on the beach.

The oystercatcher breeding season in the Richmond River area is Aug–Dec and late attempts generally are replacement clutches (Wellman *et al.* 2000). Breeding pairs were located by territory mapping *(i.e* identification of ‘pairs’ usually precedes detection of nesting attempts) and prior knowledge (known breeding territories from preceding years). The frequency for oystercatcher monitoring was mostly between 1–3 days per week. Major results are counts of breeding pairs and young fledged. This study examined breeding population size (the number of breeding pairs) and not the number of mature adults (IUCN Standards and Petitions Committee 2019), which is a less-focussed definition of population size. Reproductive rate (breeding success) was quantified as mean fledglings per breeding pair.

Pipi abundance data were obtained from several sources (Table 2). Pipis typically aggregate in discontinuous ‘bands’ that run parallel to the shoreline and, to be consistent with the majority of historical data, where quadrat-based sampling was used, abundances are reported as pipi band mean density (pipis/m^2^) in this study. Biennial counts from 2016 onwards have used a ‘feet digging’ swash zone sampling method that is more efficient than quadrat sampling. Mean feet digging pipi counts were converted to ‘pseudo pipi band densities’ following Totterman (2019a; density = 1.7 + 0.8 x mean count).

### Statistical analysis

Repeated data collection over time often leads to serial autocorrelation, where an observation at some point in time is related to, and not independent of, nearby values. The statistical modelling strategy used was: 1) fit a basic model assuming independent residuals, 2) perform a Durbin-Watson test for first-order serial correlation in the model residuals, 3) fit a generalised least squares (GLS) model with serial correlation in the residuals if required by 2). GLS adjusts the standard errors of ordinary least squares (OLS) parameter estimates for more reliable statistical inference. A continuous autoregressive GLS correlation structure was used to accommodate temporal gaps in the data. Durbin-Watson tests were limited to contiguous series within single sites. The threshold for Durbin-Watson tests was increased to *P* < 0.1 for greater statistical power for short series.

An analysis of covariance (ANCOVA) model was fitted to breeding pair counts:

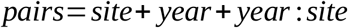

where *year:site* is an interaction term. This model fits separate linear trends for individual sites and assumes a common serial autocorrelation structure. An OLS model was assumed, rather than a generalised linear model (GLM) for counts, because trends were approximately linear over time and because modelling serial autocorrelation was relatively simple using GLS. OLS/GLS assumptions were checked with common residuals diagnostic plots. The adequacy of ANCOVA models was evaluated with likelihood ratio tests: GLS models were refitted by maximizing log-likelihood and compared to null models with a common slope (*i.e.* no *year:site* interaction; Wauchope *et al.* 2020).

The breeding population size ANCOVA for Bombing Range and South Ballina was effectively a control-impact comparison of trends. The major difference between Bombing Range (control) and South Ballina (impact) was the absence of human recreation at the former site.

With only two or three breeding pairs for Airforce and Black Rocks and most often zero at Ten Mile, those sites were too small for estimation of population size trends *(e.g.* a change of one is large relative to a population size of two or three pairs).

There was little evidence for correlated reproductive rates among sites (Spearman *r* = −0.49 to +0.31 for annual mean reproductive rates paired by year) and so unpaired comparisons were performed. There was also little evidence for temporal trends in mean reproductive rates. A simple ANOVA-like GLM was fitted to fledgling counts:

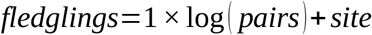

where the coefficient of one for *pairs* indicates that it is an offset (*i.e.* log(*fledglings/pairs*) = *log*(*reproductive rate*)). A quasipoisson error distribution was assumed, based on overdispersion statistics, the mean-variance relationship for residuals (Ver Hoef & Boveng 2007) and residuals diagnostic plots. GLM diagnostics used Dunn-Smyth randomised quantile residuals (Dunn & Smyth 1996). Analysis of Deviance (ANOVA for GLMs) compared the fitted ANOVA models to null models with a common mean (*i.e.* with no *site* effect). *Post-hoc* comparisons use control-impact contrasts (Dunnett’s contrasts) with Bombing Range as the control.

Some earlier studies (Owner & Rohweder 2003, Harrison 2009) have measured pipi abundance with small sample sizes and then reported means without any estimates of precision. This study reports mean pipi densities with bootstrap 95% confidence intervals. For sample sizes ≥ 10, non-parametric bootstrapping was used and bootstrap-*t* intervals are reported. For sample sizes < 10, parametric bootstrapping was used and ‘expanded percentile intervals’ (adjusted for ‘narrowness’ bias in small samples) are reported (Hesterberg 2015). Parametric bootstrapping assumed a negative binomial model for mean pipi band counts (*e.g.* mean pipis/quadrat x quadrats/band) with overdispersion parameter (‘size’) = 1.8 (Totterman 2019a). Pipi band sampling effort in Owner and Rohweder (2003; 10 x 0.0625 m^2^ quadrats) was similar to that in Totterman (2019a; 7 x 0.1 m^2^ quadrats) and so model parameters for those studies would have been similar. Sampling effort was lower in Harrison (2009; 3 x 0.11 m^2^ quadrats) and pipi band counts from that study could have been more variable than was assumed for the parametric model. Bootstrap distributions were computed with 10000 resamples.

All statistical analyses were performed using the software R version 3.5-2 (R Core Team 2018). Durbin-Watson tests for first-order autocorrelation were computed using the R package *lmtest* version 0.9-36 (Zeileis & Hothorn 2002). GLS models were fitted using *nlme* version 3.1-137 (Pinheiro *et al.* 2018). Confidence intervals for model coefficients and post-hoc comparisons were computed using *emmeans* version 1.5.3 (Lenth 2020). Randomized quantile residuals were computed using *statmod* version 1.4.32 (Dunn & Smyth 1996). Bootstrap resampling was preformed using the R package *resample* version 0.4 (Hesterberg 2015). Graphics were prepared using *ggplot2* version 3.1.0 (Wickham 2016).

## RESULTS

Historical counts of 15–16 oystercatcher breeding pairs in 1994–1996 for South Ballina, before conservation management commenced, were nearly double the recent 7–9 pairs in 2016–2018 (Table 2, Figure 2a1). A negative fluctuation from 14 to 5 pairs in 2006–2009 occurred during the 2003–2009 pipi ‘crash’. The temporary increase to 12 breeding pairs in 2010 occurred during an unusual ‘moon pipi’ *Mactra contraria* mortality event that was noted in the 2011 Pied Oystercatcher Protection Program report. Moon pipis live in the subtidal depths and are usually not available to oystercatchers.

**Figure 2.**
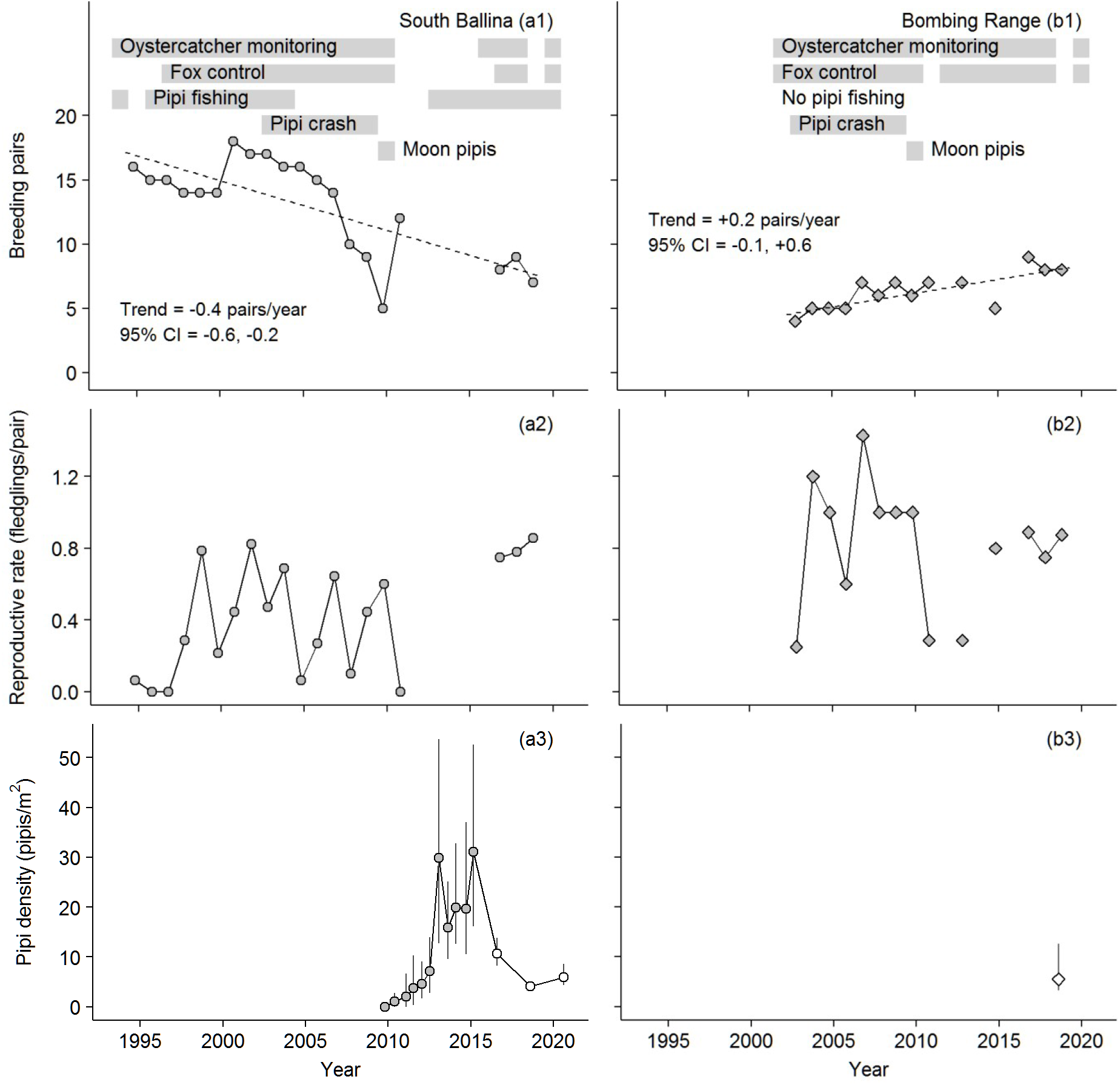
Oystercatcher breeding population size, mean reproductive rate and mean pipi density data for South Ballina (a) and Bombing Range (b). The time scale is continuous and breeding results (rows one and two) are plotted at mid-Oct, in the middle of the breeding season. The linear trends in row one are from a GLS model with beach as a covariate for year (first-order serial correlation estimate = 0.6, 95% CI = 0.2–0.9). Row three plots high confidence pipi data. Figure 3 plots all pipi data with sample sizes. Grey-filled points in rows three and four are quadrat sampling estimates, white-filled points are feet digging estimates, vertical lines are bootstrap 95% confidence intervals. The likelihood ratio test statistic for the GLS ANCOVA model, in comparison to a null model with a common slope, was 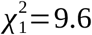 (*P* = 0.002).

There were > 207 oystercatchers fledged from beaches in the Richmond River area in 1997–2018. This number would increase if the 2011–2015 counts were complete. Despite this impressive result, the Richmond River Area beach-resident population size has decreased from 25–26 pairs in 2005–2006 to 22–23 pairs in 2016–2018 (−0.04 pairs/year). There was insufficient data for statistical evaluation of this near zero rate.

The South Ballina population size has generally decreased at −0.4 pairs/year (Figure 2a1) versus Bombing Range has increased at +0.2 pairs/year (Figure 2b1) (*year:site* interaction *P* = 0.005). Despite the response being a count variable, GLS model residuals diagnostics were satisfactory. For South Ballina and Airforce combined, the population size has decreased at −0.5 oystercatcher pairs/year (Figure 3a1) while the combined Bombing Range, Black Rocks and Ten Mile has increased at +0.3 pairs/year (Figure 3b1) (*year:site* interaction *P* = 0.008).

**Figure 3.**
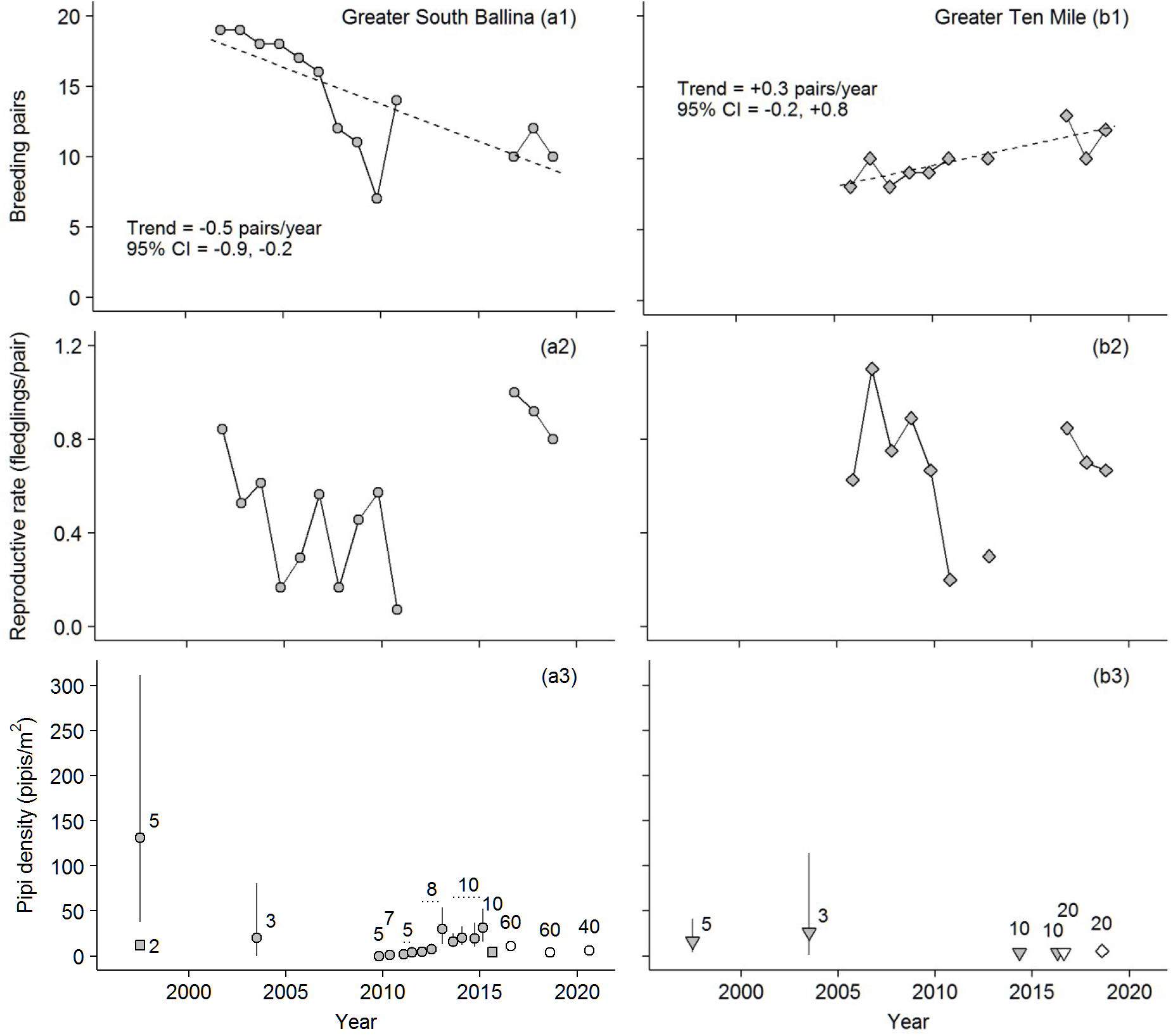
Oystercatcher breeding population size, mean reproductive rate and mean pipi density data for Greater South Ballina (South Ballina and Airforce) (a) and Greater Ten Mile (Bombing Range, Black Rocks and Ten Mile) (b). The time scale is continuous and breeding results (rows one and two) are plotted at mid-Oct, in the middle of the breeding season. The linear trends (dashed lines) in row one are from a GLS model with beach as a covariate for year (first-order serial correlation estimate = 0.5, 95% CI = 0.1–0.9). Row three plots all pipi data (South Ballina plotted as circles, Airforce as squares, Bombing Range as diamonds and Ten Mile as inverted triangles). Grey-filled points in row three are quadrat sampling results, white-filled points are feet digging results, vertical lines are 95% confidence intervals, numbers are sample sizes and dotted horizontal lines span series with constant sample sizes. It was not appropriate to estimate a confidence interval for the *n* = 2 transects for Airforce in 1997 (a3). The likelihood ratio test statistic for the GLS ANCOVA model, in comparison to a null model with a common slope, was 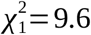 (*P* = 0.002).

The highest overall mean reproductive rate was 0.8 fledglings/pair for Bombing Range, with no public recreation, and the lowest were 0.5 fledglings/pair for South Ballina (1997–2018), with beach driving, and 0.4 fledglings/pair for Black Rocks, adjacent to a campground (Figure 4a). The large variance and high 0.7 fledglings/pair for Airforce resulted from the very small two to three pairs population size. For example, with two pairs and an average clutch size of two, there are only five possible mean fledglings per pair results (0/2 = 0.0, 1/2 = 0.5, 2/2 = 1.0, 3/2 = 1.5 and 4/2 = 2.0) and only in the worst case of zero fledglings is the mean < 0.5. For Bombing Range, Black Rocks and Ten Mile combined, the overall mean reproductive rate was 0.7 fledglings/pair and larger than 0.5 fledglings/pair for South Ballina and Airforce combined, although sample sizes were small (Figure 4b).

**Figure 4.**
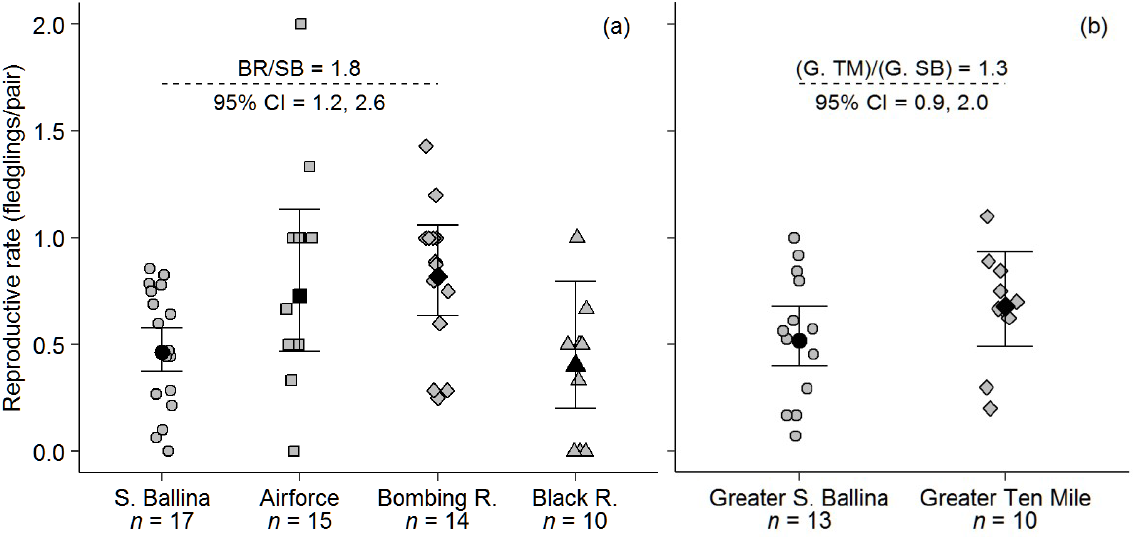
Oystercatcher mean reproductive rates for 1997–2018 (fox control commenced in 1997 at South Ballina). Ten Mile is not included in (a) because the sample size for that site was very small (*n* = 3). Greater South Ballina is South Ballina and Airforce and Greater Ten Mile is Bombing Range, Black Rocks and Ten Mile (b). Grey points are annual means and are plotted with a small amount of random horizontal ‘jitter’. Black points are overall means and error bars are 95% confidence intervals from GLMs. Dashed, horizontal lines and associated statistics are treatment versus control (Bombing Range) contrasts. With a GLM log-link, these back-transformed differences are ratios. The ‘Analysis of Deviance’ statistic for (a) was *F*_3,52_ = 4.3 (*P* = 0.009), first-order serial correlation in the GLM residuals was > −0.3 and the Durbin-Watson statistic was ≤ 2.4 (*P* ≥ 0.24). The Analysis of Deviance statistic for (b) was *F*_1,21_ = 1.5 (*P* = 0.23), first-order serial correlation in the GLM residuals was ≥ −0.1 and the Durbin-Watson statistic was ≤ 1.6 (*P* ≥ 0.71).

Small sample sizes and high variance resulted in low confidence for pre-2009 mean pipi densities (Figures 3a3, 3b3). Excluding those results, mean pipi density for South Ballina decreased from 16–31 pipis/m^2^ in 2013-2015 to 4–11 pipis/m^2^ in 2016–2020 (Figure 2a3). Pipi abundances at other sites were similar: 4 pipis/m^2^ for Airforce in 2015, 6 pipis/m^2^ for Bombing Range in 2018 and 4 pipis/m^2^ for Ten Mile in 2014–2016 (Figures 3a3, 3b3). The paired breeding results and pipi density sample size was small (*n* = 6, excluding pre-2009 mean pipi densities) and correlations for breeding pair density (pairs/km) or reproductive rate and pipi density were scattered (Figures 5a,b). There was no correlation for breeding pair density and reproductive rate (Figure 5c).

**Figure 5.**
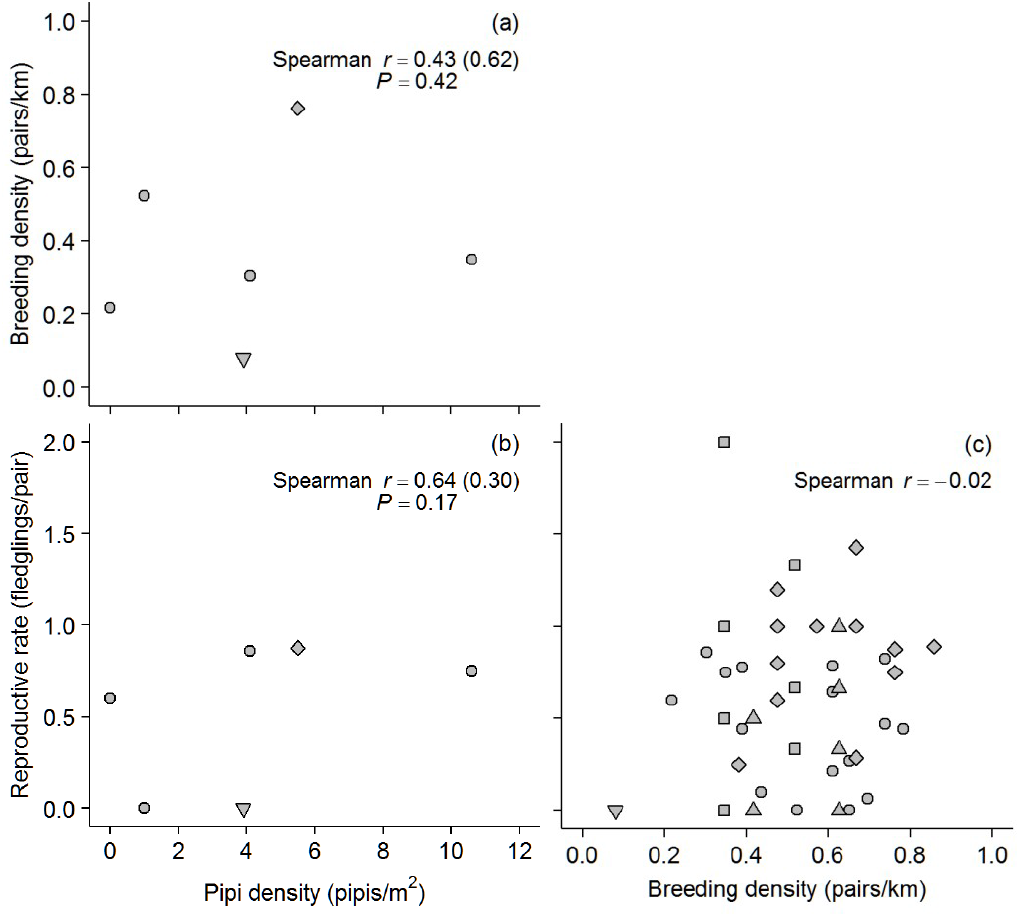
Oystercatcher breeding and pipi abundance scatterplots for all five beaches (South Ballina plotted as circles, Bombing Range as diamonds, Airforce as squares, Black Rocks as triangles and Ten Mile as inverted triangles). Low confidence mean pipi densities from 1997 and 2003 (Figures 3a4,b4) are not plotted. Correlation coefficients in parentheses refer to the complete pipi abundance data. With only two contiguous observations (2009–2010) for (a) and (b), independent observations (within variables) were assumed and *P*-values are reported.

There were six reported vehicle strike oystercatcher mortalities for South Ballina during 2001–2011 and 2015–2018. Assuming a typical six months of monitoring (Jul–Dec), at three days a week, the known 0.4 mortalities/year can be extrapolated to 2 mortalities/year, including 0.7 breeding adult mortalities/year. It is estimated that a total of 15 breeding adults have died from vehicle strikes for South Ballina in 1997–2018.

## DISCUSSION

With 22 pairs in 2018, the Richmond River Area was estimated to hold 41% of the breeding population of beach-resident Australian Pied Oystercatchers in NSW. The 48% decrease in the South Ballina population in 1994–2018 should be a serious conservation concern. The Bombing Range population did increase in 2002–2018 but could be limited by territorial behaviour at the recent 8 or 9 pairs. The maximum oystercatcher breeding pair density for South Ballina was 0.8 pairs/km in 2000 or 0.9 pairs/km for 18.7 km of available habitat (discounting 4.3 km of beach surrounding the hamlet of Patch’s Beach where there have been zero recorded nests). The maximum density for Bombing Range was 0.9 pairs/km in 2016 and was most recently 0.8 pairs/km in 2018.

The Richmond River Area oystercatcher breeding population size did not increase in 1997–2018 despite fox control successfully reducing depredation of eggs and chicks and > 207 young fledged. The majority of these fledglings cannot be accounted for. Counts of oystercatchers on NSW ocean beaches have not increased in the past two decades (Totterman 2020). There have been reports of > 100 oystercatchers in the large estuary of Port Stephens, on the Mid-North Coast of NSW, including a few individually-marked birds from the Richmond River Area, but with insufficient historical data to determine if that population has increased in the past two decades as well as few breeding records (Stuart 2010, 2011). These results do not support the hypothesis that improved breeding success will increase the oystercatcher breeding population size and a complete review of all oystercatcher breeding sites in NSW is recommended. It is predicted that, like in this study, there will be little evidence for increasing oystercatcher breeding population sizes during conservation management. Many oystercatcher fledglings have been individually marked with bands and/or leg flags over the past two decades (as part of private banding projects registered with the Australian Bird and Bat Banding Scheme) and a study of recapture and resighting data is also recommended to better understand the survival and dispersion of these birds.

The approximately linear decrease in the South Ballina oystercatcher breeding population size suggests habitat loss (degradation), *i.e.* for territorial species the population size will be reduced in proportion to the amount of habitat lost (IUCN Standards and Petitions Committee 2019). The simultaneous increase in the Bombing Range population suggests a preference for that site. Anecdotal reports indicate that vehicle-based recreation on South Ballina has increased strongly in the past decade (NSW DPIE unpub. reports 2002–2019, pers. obs.). Recent shorebird protection program reports have explicitly recommended that South Ballina be closed to 4WDs during the oystercatcher breeding season. Recent data indicate that Bombing Range does not have a higher pipi abundance than does South Ballina. Shorebird protection program reports indicate that canid nest predators are common at both sites, although South Ballina, adjoining an agricultural landscape, may have a higher proportion of foxes compared to Bombing Range, which could have a higher proportion of wild dogs (NSW DPIE unpub. reports 2002–2019). On the other hand, domestic dogs amplify human recreation disturbance for shorebirds (Glover *et al.* 2011) and are widespread along South Ballina, where dogs are commonly brought to the beach in vehicles (Totterman 2021). These comparisons indicate that the major difference between Bombing Range (control) and South Ballina (impact) is the absence of human recreation at the former site. The opposing breeding population size trends for these two sites supports the hypothesis that oystercatcher breeding population sizes are negatively impacted by human recreation disturbance. Frequent disturbance can make otherwise suitable breeding habitat less attractive, resulting in poor recruitment and a reduction in the breeding population size when old breeders die and are not replaced.

South Ballina, with uncontrolled beach driving, and Black Rocks, adjacent to a camp ground, had the lowest mean reproductive rate in this study. Bombing Range, with no public access, had the highest reproductive rates. These results support the hypothesis that oystercatcher breeding success is lower at sites with more intense human recreation disturbance. However, low breeding success is normal for oystercatchers and mean reproductive rates of 0.4–0.8 fledglings/pair from the Richmond River area are within the range 0.2–0.9 fledglings/pair from other studies for Australian Pied Oystercatchers reviewed in Taylor *et al.* (2014).

Taylor *et al.* (2014) suggested that adult oystercatcher annual survival is likely to exceed 90%. The estimated 0.7 breeding adults lost to vehicle strikes for South Ballina each year in this study suggests annual survival < 96% (= 1 – 0.7/16; assuming the 2016–2018 mean breeding population size of 8 pairs). The mortality rate could actually have increased with vehicle traffic, however the surplus of non-territorial adults at South Ballina (Totterman 2018, 2020) indicates that mortality is not limiting the breeding population size.

Aggregated along-shore distributions result in approximately quadratic mean-variance relationships for pipi abundance (Totterman 2019a). Owner and Rohweder (2003) and Harrison (2009) reported high mean pipi densities based on very small sample sizes and estimated confidence intervals in this study demonstrate that those results were hopelessly imprecise. The exceptionally high 131 pipis/m^2^ in Owner and Rohweder (2003) was also inflated by an unspecified proportion of pipi recruits < 20 mm (Owner 1997). Therefore, low quality 1997 and 2003 pipi abundance results cannot be used to infer that pipi abundance in the Richmond River area has decreased over the past two decades. More rigorous pipi sampling designs and more efficient sampling methods are recommended (Totterman 2019a,b).

With small sample sizes and a limited pipi density range, correlations for oystercatcher breeding population size and reproductive rate with pipi abundance were inconclusive. Totterman (2018) did report that oystercatcher counts increase with pipi abundance for South Ballina in 2009–2017, but cautioned that was a dynamic period, following the 2003–2009 pipi crash, and that there wasn’t a strong increase in breeding pairs. Totterman (2020) reported that habitat variables, including pipi abundance, were poor predictors for high oystercatcher territory densities. Knowing that depredation of eggs and chicks has a large effect on breeding success (Wellman *et al.* 2000), a strong correlation between oystercatcher reproductive rate and pipi abundance is *a priori* unlikely.

Conservation of resident shorebirds has commonly focussed on breeding success for reasons such as: breeding success is one of the more directly-manageable factors influencing extinction risk; positive results can be achieved quickly (*i.e.* counts of young fledged); annual reporting of breeding success results suits annual funding cycles; the idea that low breeding success leads to reductions in the population size is intuitive, and; poisoning foxes is socially acceptable to many Australians. However, control-impact comparisons in this study indicate that habitat degradation and loss resulting from human recreation disturbance can be a greater conservation threat than is depredation of eggs and chicks by foxes.

A final concern is that politics can guide conservation investments, from legislation through to funding. Politicians and Governments have apparently decided that strong protection of beach-nesting bird habitat (which involves limiting human development, recreation and banning beach driving) is less socially acceptable than controlling introduced predators and other breeding success conservation actions. Accordingly, Sch. 4 of the NSW Biodiversity Conservation Act 2016 (NSW Government 2016) does not recognise human recreation disturbance as a Key Threatening Process. The Victorian Flora and Fauna Guarantee Act 1988 (Victorian Department of Environment, Land, Water & Planning 2016) also does not list human recreation disturbance as a potentially threatening process.

## ACKNOWLEDGEMENTS

This paper was submitted to Stilt, peer-reviewed and then ‘dropped’ silently, having not been published to-date.

## APPENDIX

Richmond River Area Pied Oystercatcher Protection Program 2001–2013 and Shorebird Protection Program 2014–2018 reports were released according to the NSW Government Information (Public Access) Act 2009 (references OEH 19-452 and DPIE 20-1043).

**BT ABUS Consulting.** 2002. South Ballina Pied Oystercatcher Protection Program. 2001 Breeding Season Report. Unpub. report for the NSW National Parks and Wildlife Service.

**BT ABUS Consulting.** 2003. South Ballina Pied Oystercatcher Protection Program. 2002 Breeding Season Report. Unpub. report for the NSW National Parks and Wildlife Service.

**BT ABUS Consulting.** 2004. Richmond River Area Pied Oystercatcher Protection Program. 2003 Breeding Season Report. Unpub. report for the NSW National Parks and Wildlife Service.

**ASPECT *north.*** 2005. Pied Oystercatcher Protection Program. Report prepared following the 2004 breeding season. Unpub. report for the Richmond River Area, Northern Rivers Region, Parks and Wildlife Division, Department of Environment and Conservation.

**ASPECT *north.*** 2006. Pied Oystercatcher Protection Program. Report prepared following the 2005 breeding season. Unpub. report for the Richmond River Area, Northern Rivers Region, Parks and Wildlife Division, Department of Environment and Conservation.

**LandPartners Ltd.** 2007. Pied Oystercatcher Protection Program. Report prepared following the 2006 breeding season. Unpub. report for the Richmond River Area, Northern Rivers Region, Parks and Wildlife Division, Department of Environment and Conservation.

**Bob Moffatt EnviroServices Pty. Ltd.** 2008. Pied Oystercatcher Protection Program. Report prepared following the 2007 breeding season. Unpub. report for the Richmond River Area, Northern Rivers Region, Parks and Wildlife Division, Department of Environment and Conservation.

**Bob Moffatt EnviroServices Pty. Ltd.** 2009. Pied Oystercatcher Protection Program. Report prepared following the 2008 breeding season. Unpub. report for the Richmond River Area, Northern Rivers Region, Parks and Wildlife Division, Department of Environment and Conservation.

**Bob Moffatt EnviroServices Pty. Ltd.** 2010. Pied Oystercatcher Protection Program. Report prepared following the 2009 breeding season. Unpub. report for the Richmond River Area, Northern Rivers Region, Parks and Wildlife Division, Department of Environment, Climate Change and Water.

**Bob Moffatt EnviroServices Pty. Ltd.** 2011. Monitoring of Breeding Shorebirds and Impacts of Predators. Report prepared following the 2010 breeding season. Unpub. report for the Richmond River Area, Northern Rivers Region, Parks and Wildlife Division, Department of Environment, Climate Change and Water.

**Bob Moffatt EnviroServices Pty. Ltd.** 2013. Pied Oystercatcher Protection Program. Summary report prepared following the 2012 breeding season. Unpub. report for the Richmond River Area, Northern Rivers Region, NSW National Parks and Wildlife Service, Office of Environment and Heritage.

**Naturecall Environmental.** 2014. Shorebird Monitoring, North NSW Coast. Unpub. report for the NSW National Parks and Wildlife Service, Office of Environment and Heritage.

**Reconeco Pty. Ltd.** 2015. Richmond River Area Shorebird Protection Program 2014. Summary Report for the 2014 Breeding Season. Unpub. report for the Richmond River Area, Northern Rivers Region, NSW National Parks and Wildlife Service.

**Naturecall Environmental.** 2016. Vertebrate Pest Management and Shorebird Monitoring Report, North NSW Coast. Unpub. report for the NSW National Parks and Wildlife Service, Office of Environment and Heritage.

**Reconeco Pty. Ltd.** 2017. Richmond River Area Shorebird Protection Program 2016. Summary Report for the 2016/17 Breeding Season. Unpub. report for the Richmond River Area, Northern Rivers Region, NSW National Parks and Wildlife Service.

**Reconeco Pty. Ltd.** 2018. Richmond River Area Shorebird Protection Program 2017. Summary Report for the 2017/18 Breeding Season. Unpub. report for the Richmond River Area, Northern Rivers Region, NSW National Parks and Wildlife Service.

**Reconeco Pty. Ltd.** 2019. Richmond River Area Shorebird Protection Program 2018. Summary Report for the 2018/19 Breeding Season. Unpub. report for the Richmond River Area, Northern Rivers Region, NSW National Parks and Wildlife Service.

## Notes

### Competing Interest Statement

The authors have declared no competing interest.

### Summary of Updates

Still not published! I made a few minor changes/corrections and consider this to be the final report.

